# Not just the sum of its parts: geographic variation and non-additive effects of pyrazines in the chemical defence of an aposematic moth

**DOI:** 10.1101/2022.05.28.493811

**Authors:** Cristina Ottocento, Anne E. Winters, Bibiana Rojas, Johanna Mappes, Emily Burdfield-Steel

## Abstract

Chemical defences often vary within and between populations both in quantity and quality, which is puzzling if prey survival is dependent on the strength of the defence. We investigated the within-and between-population variability in chemical defence of the wood tiger moth (*Arctia plantaginis*). The major components of its defences, SBMP (2-sec-butyl-3-methoxypyrazine) and IBMP (2-isobutyl-3-methoxypyrazine) are volatiles that deter bird attacks. We expected the variation to reflect populations’ predation pressures and early-life conditions. To understand the role of the methoxypyrazines, we experimentally manipulated synthetic SBMP and IBMP and tested the birds’ reactions. We found a considerable variation in methoxypyrazine amounts and composition, both from wild-caught and laboratory-raised male moths. In agreement with the “cost of defence” hypothesis, the moths raised in the laboratory had a higher amount of pyrazines. We found that SBMP is more effective at higher concentrations and that IBMP is more effective only in combination with SBMP and at lower concentrations. Our results fit findings from the wild: the amount of SBMP was higher in the populations with higher predation pressure. Altogether, this suggests that, regarding pyrazine concentration, more is not always better, and highlights the importance of testing the efficacy of chemical defence and its components with relevant predators, rather than relying only on results from chemical analyses.

## 1. Introduction

Aposematism is an anti-predatory strategy in which organisms present primary defences (warning signals, which operate before the attack, see Ruxton et al. 2004; Rojas et al., 2015) and secondary defences (e.g., chemical defences, which operate upon attack, see Ruxton et al., 2018). Chemical defences are one of the most common types of secondary defences used by aposematic species (Eisner & Siegler, 2005; Speed et al., 2012). Aposematic colouration can vary greatly among individuals of the same population (Speed et al., 2012; Briolat et al. 2019). This may seem contradictory at first, as the survival of aposematic individuals relies mostly on the predator’s ability to detect and memorise a certain colour pattern associated with a disgusting taste (Mallet and Singer, 1987; Sherratt and Speed, 2004; Sherratt, 2008). There are several potential non-mutually exclusive mechanisms to explain variation in warning coloration, including opposing selection pressures, variable selection or heterozygote advantage just to name few (rev. Endler & Mappes 2004; Briolat et al. 2019). Also, variation in the composition and dose of the chemical defences requires for evolutionary explanation because prey survival depends on the strength of the defence (Speed et al., 2012). Despite this, variation in chemical defences both between-and within-populations exists (in ladybirds *Harmonia axyridis*, see Bezzerides et al., 2007 and Arenas et al., 2015; in poison frogs, *Dendrobates tinctorius*, see Lawrence et al., 2019; in nudibranchs *Goniobranchus splendidus*, see Winters et al., 2019; in Heliconiini butterflies, see Sculfort et al., 2020). Between-population variation may reflect the evolutionary history of populations; for example due to differences in predation pressure, the environmental conditions and availability of resources (Joron & Iwasa 2005; Sherratt, 2006; Briolat et al., 2019). Within-population variation could be explained as well by a quantitative variation in toxins due to environmental heterogeneity if the diversification in the defence reflects the nutrients available in that habitat (Brower et al., 1967; Bowers, 1992). However, the diversity in chemical defences within the same population may reflect not only the environmental conditions but also other factors such as body size, sex and age (Alonso-Mejia & Brower, 1994; Hudson et al., 2021). The variation in biochemical profiles may have an adaptive significance, especially in organisms that synthesise their toxins *de novo*. This may be due to competition for resources (due to obtaining toxins from the diet), or the effect of also using the chemical defences for communication, for example, pheromones (Pfeiffer et al., 2018; Böttinger et al., 2021).

Chemical defences in invertebrates may be associated both with a distasteful flavour and an unpleasant odour for the predators. One of the prevalent types of odorants known is pyrazine, a heterocyclic aromatic organic compound and an organoleptic agent involved in the release of a warning smell in many aposematic insects (Moore & Brown, 1981; Rothschild & Moore, 1987, Guilford et al. 1987). Pyrazine has a distinctive smell and a disgusting taste that has been shown to help predators learn the association between a warning signal and a secondary defence (Rowe & Guilford, 1996).

Here we investigated whether different populations of the wood tiger moth, *Arctia plantaginis*, differ in the composition of their chemical defences, and how wild predators respond to said defences. Previous studies (e.g. see Darst & Cummings 2006; Darst et al. 2006; Maan & Cummings 2012; Cummings & Crothers 2013) show that chemical defences are often measured, but the response of relevant predators is often overlooked (see Weldon, 2017). These measurements may thus fail to reflect what happens in the wild, which can be problematic if, for example, predator response does not correlate positively, or consistently, with the measured amounts of the defence components. For example, the predators may ignore or cannot detect the variation present in the chemical defence (Lawrence et al., 2019) or different compounds are effective only against one predator type (Rojas et al. 2017). Therefore, measuring the response(s) of relevant predators is crucial for understanding what kind of evolutionary implications chemical variation may have.

We tested whether the efficacy of chemical defences can be extrapolated from their composition, using European and Caucasian wood tiger moth populations both wild-caught and raised in the laboratory. The wood tiger moth, *Arctia plantaginis*, is a chemically defended, warningly-coloured species. The major components of this moth’s defences, the methoxypyrazines SBMP (2-sec-butyl-3-methoxypyrazine) and IBMP (2-isobutyl-3-methoxypyrazine), are synthesised *de novo* and secreted as reflex blood in response to attacks by avian predators. First, we investigated within-and between-population variation in the amount and composition of methoxypyrazines from wood tiger moths collected in Estonia, Finland, Scotland and Georgia. We expected variation in chemical defences to reflect populations’ predation pressure, being weaker and having greater variability in populations with low predation pressure (Estonia and Finland; Rönkä et al. 2020) and stronger with less variability in populations with high predation pressure (Scotland and Georgia; Rönkä et al. 2020). To understand the role of each of the two methoxypyrazines, we experimentally manipulated the strength of pure (synthetic) SBMP and IBMP, both separately and combined, and tested the reactions of wild predators toward them. Finally, we compared the natural variation of wild moth chemical defences to the defence variation of moths that were raised in the laboratory in standard conditions, and then tested predator reactions toward their defensive secretions. If pyrazine production is costly and early life conditions are important for chemical defences, we expect to see less variability, higher amounts of pyrazines and stronger predator reactions toward moths raised in the laboratory *ad libitum* compared to wild moths.

## 2. Materials and methods

### 2.1 Study species

The Wood tiger moth *Arctia plantaginis* (formerly *Parasemia plantaginis*; Rönkä et al., 2016) is a diurnal aposematic species, and presents two different types of chemical fluids: one is produced from the abdomen and it is a deterrent to ants. The thoracic fluid, on the other hand, is an effective deterrent to birds (Rojas et al., 2017) thanks to two methoxypyrazines: SBMP and IBMP. These methoxypyrazines are not sequestered directly by plants but are produced *de novo* (Burdfield-Steel et al., 2018). The wood tiger moths display conspicuous hindwing colouration that varies both geographically and locally (Watson & Goodger, 1986; Chinery, 1993; Nokelainen et al., 2013) throughout the Holarctic region (Hegna et al., 2015; Leraut, 2006; Powell & Opler, 2009). While it is known for its colour polymorphism in male hindwing colouration (Hegna et al., 2015), some populations are also known to be monomorphic; for example, in Northern Scotland and in Estonia, the hindwings of adult males are either yellow or white, respectively (Hegna et al., 2015). Georgian wood tiger moths in the Caucasus region are phenotypically different to the other European populations (Hegna et al., 2015; Rönkä et al., 2016; Yen et al., 2020), as the hindwing colouration in males presents continuous variation from yellow to red (Rönkä et al., 2020). In Finland, wood tiger moths produce only one generation per year in the wild (Ojala et al. 2005, Lindstedt et al. 2010). Under greenhouse conditions, however, it is possible to obtain up to three generations per year. The larvae are polyphagous, feeding on a variety of different plants such as *Taraxacum sp*. (dandelion), *Plantago sp*., *Rumex sp*. and *Vaccinium uliginosum* (Ojala et al., 2005).

### 2.2 Measurement of pyrazine levels across populations

#### Collection of thoracic fluids

Moths from wild populations were collected between 2015 and 2018 in four countries, Finland, Estonia, Georgia, and Scotland. Moths were caught either in nets or in pheromone traps baited with laboratory-reared females. Upon capture, the moths were kept in individual containers and either had their thoracic fluids sampled the day after capture, or were transported back to the University of Jyväskylä for sampling. Laboratory-reared moths were taken from populations founded from individuals from the same countries and maintained at the University of Jyväskylä. All laboratory-reared individuals originated from eggs laid and reared at the University, although the length of time that the stock they originated from was kept in the laboratory varied. Larvae were kept at approximately 25°C during the day, dropping to 15-20°C at night. For the Finnish and Estonian populations, larvae were overwintered every third generation at 5°C during the third instar. Larvae were housed in clear plastic tubs in family groups of no more than 30, fed with *Taraxacum sps*. (dandelion) and misted with water daily. The only exception to this was the Georgian population, whose diet in the laboratory, besides *Taraxacum sp*., was supplemented with *Plantago sp*. and *Rumex sp*.. Tubs were cleaned daily as needed and uneaten food was replaced. Upon pupation, individuals were kept individually in vials at 25°C until eclosion.

In all cases, the protocol for sampling the thoracic fluids was the same and followed the method previously described in Rojas et al. (2017) and Burdfield-Steel et al. (2019). Moths were stored in chilled conditions (approximately 4 degrees) until 1 hour prior to sampling, at which point they were provided with water (either in droplets or on a damp paper towel) to rehydrate and placed at room temperature to become active. Thoracic fluids were collected by pinching just below the prothoracic section of the moths with a pair of tweezers. This stimulated the release of the defence fluid, which was then collected with 10 µl glass capillaries and the volume was measured with a calliper. Samples were then transferred to glass vials and stored at -20°C until analysis.

#### Measurement of methoxypyrazines

Prior to GC-MS analysis, samples were thawed and mixed with a 200-µl NaCl solution (3%). Measurement of the pyrazines was done following the methods of Cai et al. (2007) as described in Burdfield-Steel et al. (2018). Pyrazines were extracted from the headspace of fluid samples using SPME fibers (StableFlex 1-cm fibers with Divinylbenzene/Carboxen/Polydimethylsiloxane coating, Sigma-Aldrich, Darmstadt, Germany) for 30 min at 37°C. GC/MSD and analyses were carried out on an Agilent 6890 series GC system equipped with a Zebron ZB-5HT Inferno (Phenomenex Inc., Torrance, CA) column (length 30 m, 0.25 mm I.D. with a film thickness of 0.25 µm) connected to an Agilent 5973N MSD. Fibers were manually loaded into the injector using a splitless injection mode, and the inlet temperature was set to 260°C. Helium was used as a carrier gas at a constant flow rate of 0.8 ml/min. The oven temperature was programmed as follows: 3 min at 60°C then ramped to 170°C at a rate of 7°C/min and from 170 to 260°C at a rate of 20°C/min and kept at that temperature for an additional 5 min. SBMP and IBMP were detected using selected ion monitoring of ions 124, 138, and 151. The chromatograms and mass spectra were evaluated using Agilent Chemstation (v. G1701CA) software and the Wiley 8th edition mass spectral database and the methoxypyrazines were identified using the ratio of these detected ions from the NIST webbook page (Stein), as well as by comparison with standards of SBMP and IBMP. The amount of the two methoxypyrazines in the sample was calculated by comparison with known amounts of the standards, run in the same manner as the fluid samples.

#### Measure of methoxypyrazine statistical analysis

All statistical analyses were carried out with the software R v. 4.1.2 (R Development Core Team, 2019) using the RStudio v. 1.2.1335 interface (RStudio Team, 2018). We tested the effect of population (Finnish, Estonian, Georgia, Scotland) and rearing environment (hereon referred to as origin: wild vs. laboratory) on the amount (ng) of SBMP, IBMP, the ratio of IBMP to SBMP and the total amount of pyrazine using linear mixed-effects models with a normal distribution in the package lmer4 (Bates et al., 2015). In each model, population, ‘origin’ and the interaction between the two were set as the explanatory variables and ‘year’ was included as a random factor, to account for the non-independence of data gathered within the same year. A Watson–Williams F-test (pairwise comparison) was applied to compare variance in the amount SBMP and IBMP from wood tiger moths raised in the laboratory vs wild, and Bartlett test (Bartlett M. S., 1937) was applied to compare variance in the amount of each pyrazine across populations.

### 2.2 Pure pyrazine assay

Synthetic SBMP and IBMP (from Supelco, Sigma-Aldrich) were diluted in water at the University of Jyväskylä to the following concentrations: 0.05, 0.1, 0.5, and 1 ng/µl. A 50/50 blend of SBMP and IBMP was also made such that each dilution (0.05, 0.1, 0.5, and 1 ng/µl) was the total additive concentration of the two pyrazines combined. These dilutions were then refrigerated at ∼4°C for no more than one month before use in the experiment.

We used blue tits (*Cyanistes caeruleus*) as a model predator to test their response to the pure methoxypyrazines. This species is common in Finland, easy to capture and possible to keep in captivity for short periods of time necessary for the experiments (Rönkä et al., 2018). Furthermore, blue tits are thought to be a natural predator of wood tiger moths, they have an overlapping distribution range and have already been used in different studies on the chemical defences of wood tiger moths (Rojas et al. 2017; Rönkä et al. 2018; Burdfield-Steel et al. 2019; Rojas et al. 2019).

The birds used for the experiment were caught at Konnevesi Research Station (Central Finland), from January to March in the years 2017 - 2019, maintained individually in plywood cages with a perch, water bowl and *ad libitum* food, and kept on a 12:12 h light:dark cycle. Each bird was weighed before and after the experiment, and ringed, sexed and aged before being released to the place of capture. Birds were used with permission from the Central Finland Centre for Economic Development, Transport and Environment and licensed by the National Animal Experiment Board (ESAVI/9114/04.10.07/2014) and the Central Finland Regional Environment Centre (VARELY/294/2015). All experimental birds were used according to the ASAB/ABS Guidelines for the treatment of animals in behavioural research and teaching (Association for the Study of Animal Behaviour, 2020). Once trapped from a feeder (13 × 17 × 40 cm box) with peanuts as a bait (Ham et al. 2006; Lindstedt et al. 2011; Nokelainen et al. 2012), the birds were housed individually, and then transferred to the experimental boxes for training and habituation purposes. Before starting the trials, the birds were familiarised with the experimental boxes and trained to eat oats. The experimental box was made from plywood (50 × 60 × 45 cm) and lit with a light bulb showing the entire visible daylight spectrum (Nokelainen et al. 2012). Inside the box, there was a water bowl, a perch and a small barrier that allowed observers to clearly see when the birds first noticed the food item presented. This barrier was in front of a moving hatch on which the food was placed (Nokelainen et al. 2012).

A total of 79 blue tits were used to measure bird responses to pure pyrazines. Each bird was used in the experiment only once and was assigned a single treatment. Birds were first trained to eat water-soaked oats before use in the assay. Each assay consisted of five trials. In the first trial, birds were offered water-soaked oats to ensure they were motivated to feed and, in the last trial, birds were again offered water-soaked oats to discard satiation. During trials 2,3 and 4 each bird was presented with 3 oats per trial on a small white dish. Each oat was covered with 8µl of either water (as a control treatment) or one of the pure pyrazine treatments. Therefore, only trials 2, 3, and 4 are used in the analysis. In each trial we recorded hesitation time (measured as time in seconds from seeing the oat to pecking/eating the first oat), the proportion of the oats eaten (to the nearest 10%), beak cleaning (a disgust behaviour measured as the number of bouts where the bird wiped its beak against a surface such as the perch), drinking (the number of times the bird drank water, which is a behaviour that can increase in response to distasteful food), and trial duration (from the time the oats were seen by the bird until they were consumed – or max 300 seconds if some of the oats remained). All trials were video recorded using a hole at the top of the experimental enclosure.

#### Pure pyrazine assay statistical analysis

All analyses were conducted using R version 4.1.2 (R Development Core Team, 2019). To test whether bird hesitation time differed among treatments, we used a cox proportional hazards model using the package coxme (Therneau, 2020). To test whether the percentage of oats eaten or counts of bird beak cleaning and water drinking behaviours differed among treatments, we first excluded observations from birds that did not eat any of the oats and included trial duration as an offset term in the models. We then used Generalized Linear Mixed Models (GLMM) with Poisson distribution using package lme4 (Bates et al. 2015). For each bird response variable, we tested whether pyrazine treatments differed from the control treatment. In each model, the predictor variables included the chemical treatment (a categorical variable with different levels for each pyrazine and concentration) and trial number (2,3,4) to test whether birds altered their behaviour as the trials progressed. Each model also included bird age, sex, and weight as co-variates and bird ID as a random factor. Automated model selection was performed using the dredge function of the MuMIn package (Barton et al. 2015) with chemical treatment and trial number set as fixed in all models. In all cases, the simplest model within delta 2 of the top model contained only chemical treatment and trial number as fixed factors and bird ID as a random factor, and this was chosen as the final model.

### 2.3 Bird response to moths’ defensive fluid

#### Larval diet in the laboratory and wild moths

The male wood tiger moths used in this work were obtained from a laboratory stock kept at the greenhouse of the University of Jyväskylä with wild-caught individuals from Finland and Georgia. The temperature in the greenhouse was between 20°C and 30°C during the day and decreased to 15°C-20°C during the night. Daylight lasted for approximately 20 hours. After hatching, larvae were fed with lettuce and *Taraxacum sps*., Georgian larvae, as mentioned above, were also fed with *Plantago sp*. and *Rumex sp*. When the adults emerged from pupae, they were given water and stored at 4 degrees to slow their metabolic rate. Thoracic fluids were obtained in 2015 and 2017 by gently squeezing with tweezers below the thoracic section of the moth, and further collected with a 10 μl glass capillary and stored in Eppendorf tubes at -18 degrees. Prior to fluid collection, the moths were initially kept between 20 and 25 degrees for thirty minutes.

#### Predator assay

Blue tits (*C. caeruleus*; n=116) were used for the predator assay as described in the previous section (pure pyrazine analysis). We used fluids from 34 wild Finnish males, 21 laboratory Finnish males, 44 wild Georgian males and 8 laboratory Georgian males. In addition, we offered water to 9 birds which were used as controls. Because the moths had different volumes of thoracic fluid, the fluid of each individual fluid was diluted proportionally with water to reach a total volume of 15 μl of fluid. Then, the 15 µl were divided into two samples of 7 μl each, which were offered to the same bird. The same amount of water was offered to control birds. During bird training sessions, we put 4 oat flakes and 3 sunflower seeds on a small white plate. Only when the birds ate all the oats in the training phase, did the experiment begin.

Each bird experienced 4 trials, one after the other, with 5-minutes intervals. In each trial, the bird was presented with a plate containing one oat flake. We decided to use only four trials and one oat flake per plate (compared to the five trials and three oat flakes in the pure pyrazine assay) because of the availability of the amount of volume of chemical defence fluid per moth individual, following the methodology of Burdfield-Steel et al. 2019. The first and last trials were done with oats soaked in water to ensure that the bird was motivated to eat (first) and that the bird was still hungry (last). In the second and third trials, the bird was presented with an oat soaked in 7μl of the defensive fluids of the same moth. The trial ended two minutes after the bird had eaten the whole oat, or after a maximum duration of 5 minutes if the bird did not eat the whole oat. In each trial, we recorded the hesitation time (the time in seconds that occurs from the moment when the bird sees the oat to when they pecking/eating it); the proportion of oat eaten(to the nearest 10%); beak cleaning (number of times the bird wiped its beak against a surface - e.g. the perch); the drinking (number of times the bird drinks water, as a response to the distaste of the food); the trial duration (from the time the oats were seen by the bird until they were consumed – or max 300 seconds if the oat was not eaten). All trials were video recorded.

#### Predator assay statistical analysis

The statistical analyses were conducted using the software R v. 4.1.2 (R Development Core Team, 2019) using the RStudio v. 1.2.1335 interface (RStudio Team, 2018). The behaviours of the birds were first compared to a water-only control to determine if the thoracic fluid of the moths elicited an adverse reaction in the predators. To test the difference in hesitation time in response to thoracic fluids from wild and laboratory Finnish and Georgian wood tiger moths, we used a cox proportional hazards model using the package coxme (Therneau et al. 2020). The percentage of oats eaten per minute, birds’ beak cleaning and water drinking behaviours, were tested using general linear mixed-effects model with Poisson distribution using package lme4 (Bates et al. 2015). Each model included bird ID as a random factor. The population, origin (wild/laboratory) and the interaction between the two were set as fixed factors. Also, the trial with duration as an offset was included as an explanatory variable, while the hesitation time, percentage of oat eaten, the beak wiping frequency and drinking frequency were set as response variables. These models only looked at observations where the proportion eaten was greater than zero. The treatments that showed a different reaction to the water control (hesitation time, percentage of oat eaten per minute and water drinking) were then compared using Tukey–Kramer post hoc test for multiple comparisons and excluding the water control group. Statistical significance was set at p < 0.05.

## 3. Results

### 3.1 Geographic variation in pyrazines

#### Differences in pyrazines across populations

The quantity of SBMP in the thoracic fluids of wood tiger moths was significantly influenced by the interaction between population and origin (Chisq = 10.68; df = 3; Pr(>Chisq)=0.014; Table S1, Figure 1): laboratory-raised individuals from Estonia, Finland and Georgia have a higher amount of SBMP compared to the wild individuals from the same populations, whereas this trend was the opposite for the Scottish population (i.e., wild individuals from Scotland had a higher amount of SBMP than their laboratory-raised counterparts; Table S1, Figure 1). The IBMP amount found in laboratory-raised individuals was significantly higher than that of wild individuals (Chisq =11.08; df = 1; Pr(>Chisq)=0.0009; Table S2 Figure 1). Populations differed in the IBMP amount found in their thoracic fluids (Chisq = 27.25; df = 3; Pr(>Chisq)=5.224e-06; Table S2, Figure 1). We found no effect of the interaction between population and origin on the quantity of IBMP (Chisq = 5.73; df = 3; Pr(>Chisq)=0.125; Table S2, Fig 1).

**Fig.1.**
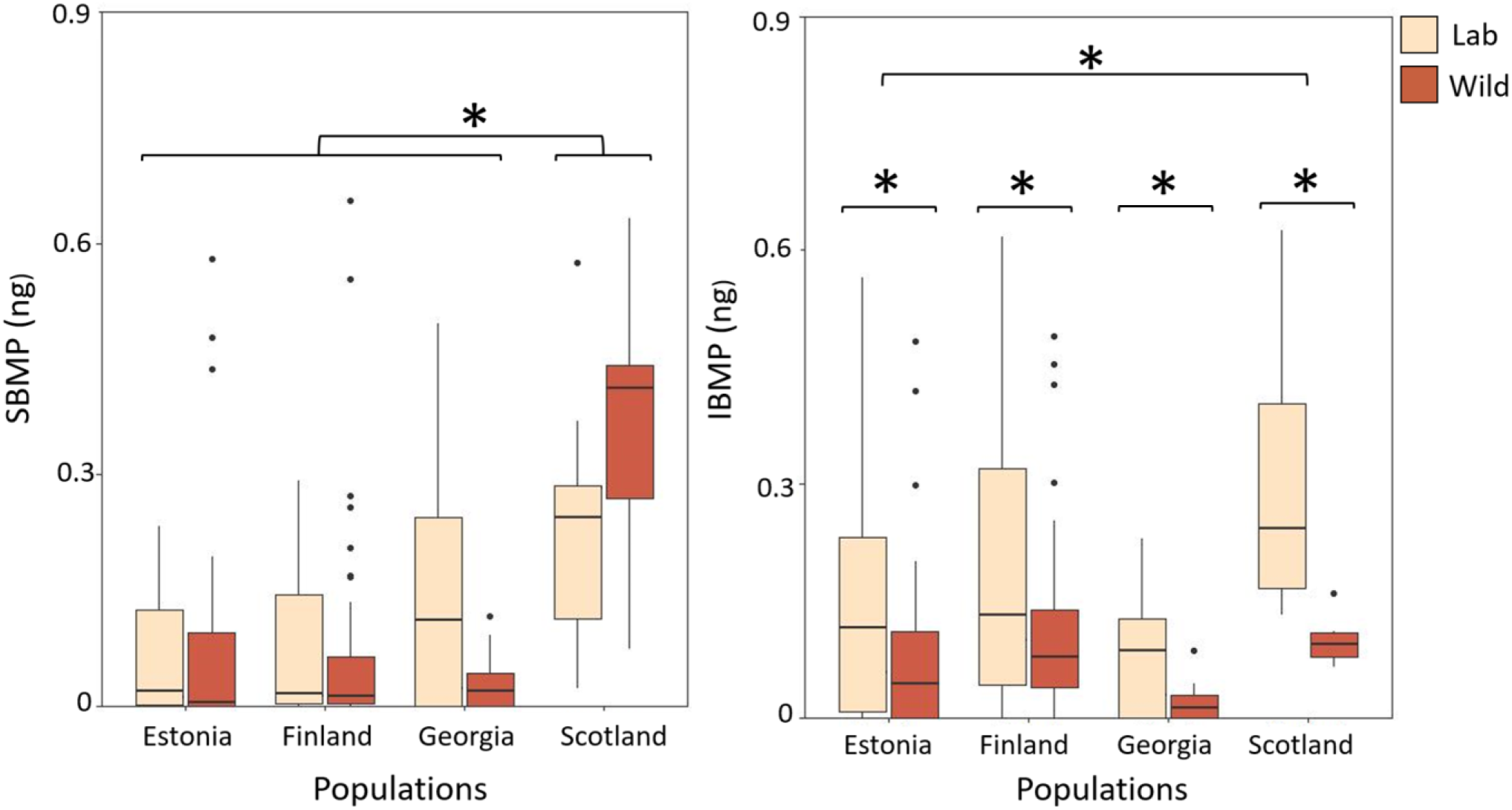
SBMP and IBMP amount in nanograms for each population (note this is the total amount of SBMP and IBMP in the thoracic fluids in nanograms, not the concentration). Boxes show the median and the 25th and 75th percentiles of data distribution. Vertical lines show the data range.

The variance between laboratory and wild wood tiger moth was equal in the amount of SBMP (F = 0.99, num df = 63, denom df = 91, *p* = 0.99; Figure 1), but not for the amount of IBMP (F = 2.70, num df = 63, denom df = 91, *p* = 1.48e-05; Figure 1): wild wood tiger moths present the lowest variance in the amount of IBMP (see table S3). Variance was different between populations in the amount of SBMP (Bartlett’s K-squared = 10.31, df = 3, *p* = 0.016) and the amount of IBMP (Bartlett’s K-squared = 16.745, df = 3, *p* = 0.0008). The Finnish wood tiger moth population presented the highest variance in SBMP amount, on the other hand, the Estonian population presented the highest IBMP variance, followed by the Finnish population (see table S4).

There were significant differences in the ratio of IBMP to SBMP between populations (Chisq =18.0482; df = 3; Pr(>Chisq)=0.0004 ***; Table S5, Figure 2), but neither origin (Chisq =2.6254; df = 1; Pr(>Chisq)=0.105; Table S5, Figure 2), nor the interaction between population and origin had an effect on the ratio of IBMP to SBMP (Chisq =1.9633; df = 1; Pr(>Chisq)=0.58; Table S5, Figure 2). The interaction between population and origin has a significant effect on the total amount of pyrazine (Chisq =10.23; df = 3; Pr(>Chisq)=0.017 *; Table S6, Figure 3

**Fig. 2.**
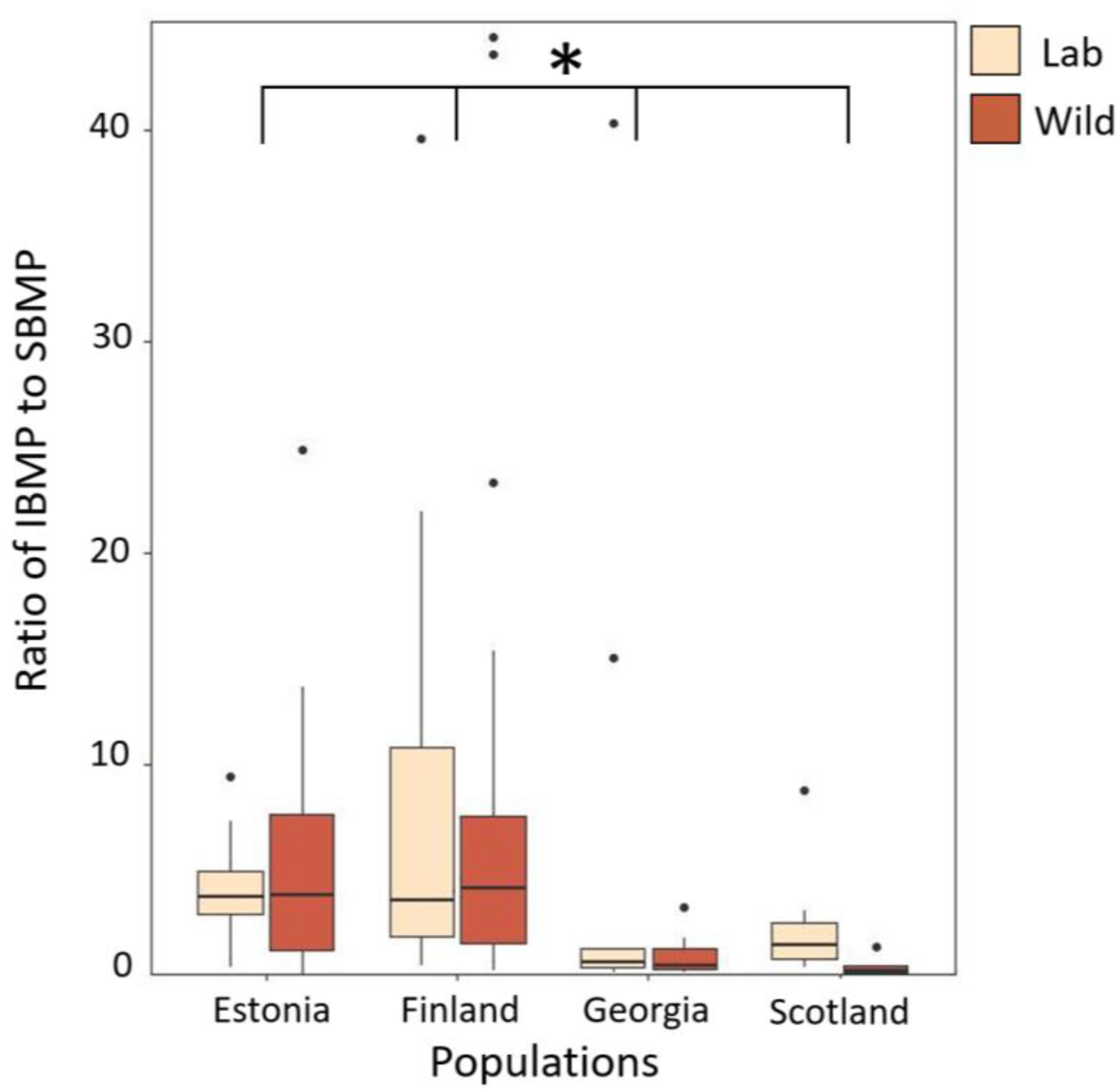
Ratio of IBMP to SBMP and total amount of pyrazine (in nanograms) for each populations. Boxes show the median and the 25th and 75th percentiles of data distribution. Vertical lines show the data range.

**Fig 3.**
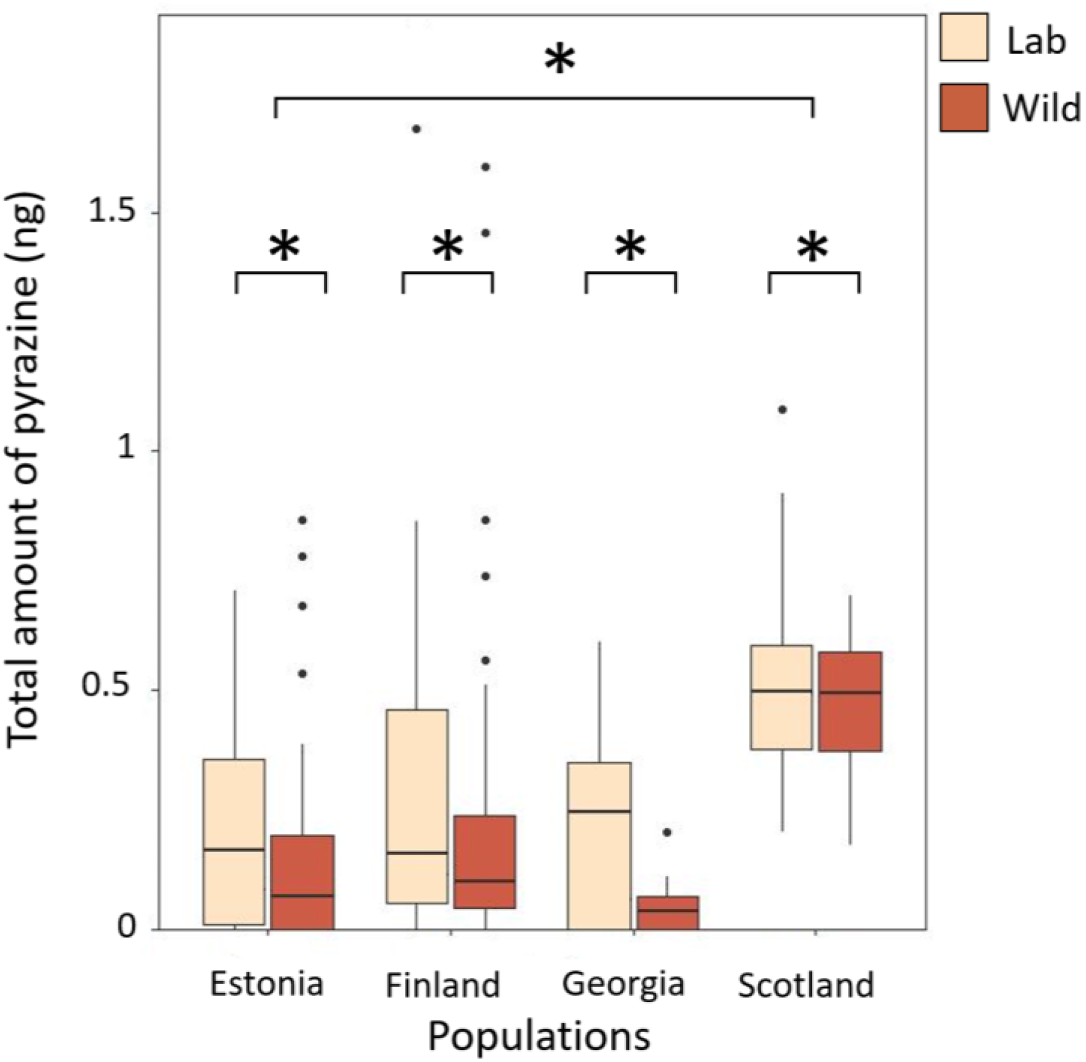
Ratio of IBMP to SBMP and total amount of pyrazine (in nanograms) for each populations. Boxes show the median and the 25th and 75th percentiles of data distribution. Vertical lines show the data range.

### 3.2. Pure pyrazine essay

Birds hesitated longer to eat oats in later trials (coef ± SE = -0.21± 0.09, z = -2.28, *p* = 0.023), but none of the pyrazine treatments differed significantly from the control (Table S7, Figure S1). However, there was a trend for birds to hesitate longer before eating oats of SBMP 0.1 ng/µl (coef ± SE = -1.33 ± 0.68, z = -1.95, *p* = 0.051) and 1.0 ng/µl (coef ± SE = -1.34 ± 0.69, z = -1.95, *p* = 0.052) concentrations compared to the control (Figure S1). Birds ate a smaller proportion of the oats in later trials (estimate ± SE = -0.07 ± 0.01, z = -5.60, *p* < 0.001, Figure 4). In addition, the proportion of oats birds ate was significantly less than the control for SBMP at the three highest concentrations: 0.1 ng/µl (estimate ± SE = -2.70± 1.28, z = -2.11, *p* = 0.04, Figure 4), 0.5 ng/µl (estimate ± SE = -4.76 ± 1.31, z = -3.64, *p* < 0.001, Figure 4), and 1.0 ng/µl (estimate ± SE = -3.01 ± 1.28, z = -2.34, *p* < 0.019, Figure 4) and for the 50/50 blend of SBMP + IBMP at the 0.05 ng/µl (estimate ± SE = -2.57 ± 1.28, z = -2.00, *p* = 0.05, Figure 4) and 1.0 ng/µl concentrations (estimate ± SE = -2.35± 1.20, z = -1.96, *p* = 0.050, Figure 4), but no concentrations of IBMP differed from the control (Table S 8, Figure 4).

**Fig. 4.**
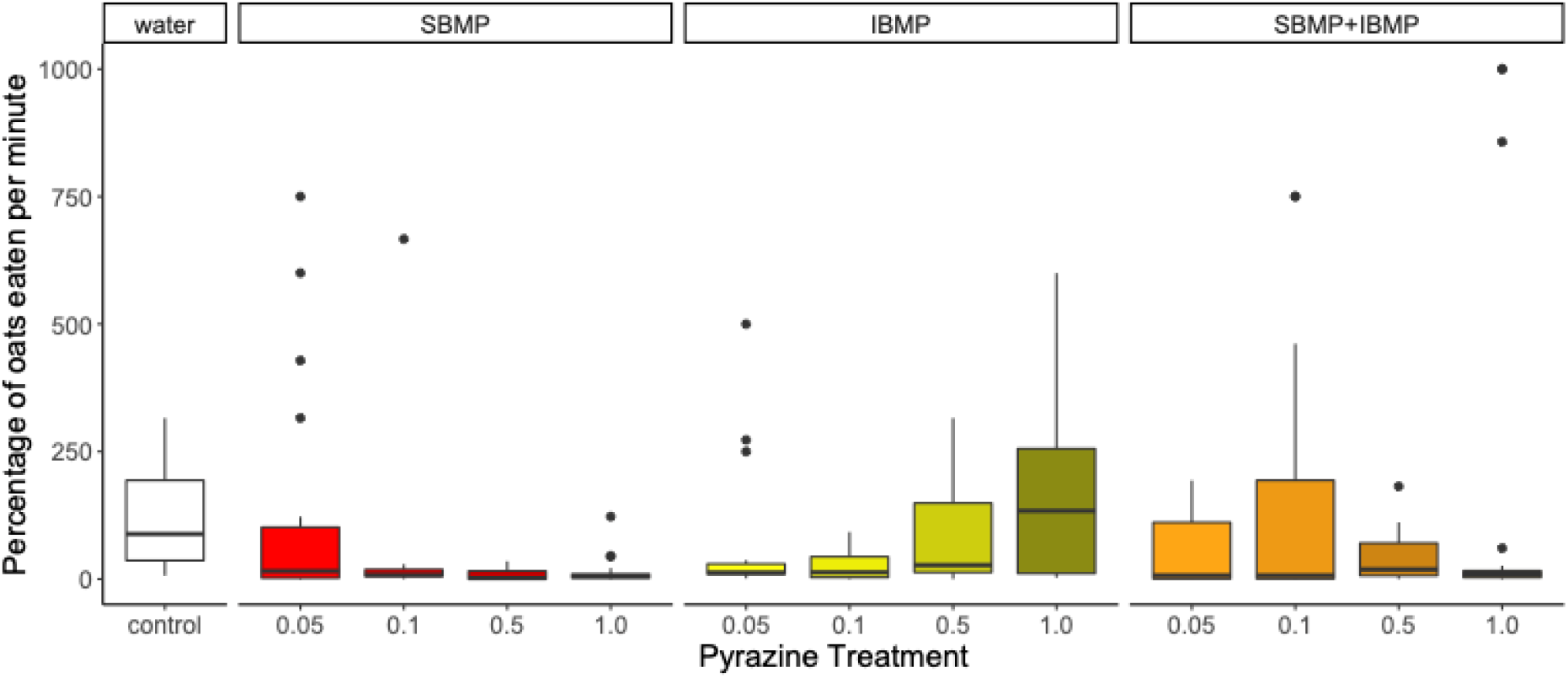
Percentage of fluid-soaked oats eaten per minute for each pyrazine type (SBMP = red, IBMP = yellow, SBMP+IBMP = orange) and ng/*u*l concentration (higher concentrations shown in darkening shades) compared to the water control. Boxes show the median and the 25th and 75th percentiles of data distribution. Vertical lines show the data range.

Bird beak wipes did not change across trials (estimate ± SE = -0.02 ± 0.05 z = -0.43, *p* = 0.668), and none of the pyrazine treatments differed significantly from the control (Table S9, Figure S2). However, there was a trend for birds to wipe their beaks more after eating oats of the SBMP+IBMP 0.5 ng/µl concentration compared to the control (estimate ± SE = 1.58 ± 0.82, z = 1.93, *p* = 0.054, Figure S2).

Birds drank more water in later trials (estimate ± SE = 0.34 ± 0.07, z = 4.60, *p* < 0.001). In addition, birds drank more water in response to the SBMP+IBMP 0.5 ng/µl concentration compared to the control (estimate ± SE = 3.06 ± 1.41, z = 2.17, *p* = 0.030, Figure S3). There was also a trend for birds to drink more water in response to the IBMP 0.05 ng/µl concentration compared to the control (estimate ± SE = 2.46 ± 1.42, z = 1.74, *p* = 0.083, Figure S3), but no concentrations of SBMP differed from the control (Table S10, Figure S3).

### 3.3 Bird response to moths’ defensive fluid

Following the measurement of the pyrazine levels across populations and the bioassay testing the response of wild-caught predators to the pure pyrazine, we tested bird response to the thoracic fluid of Finnish and Georgian laboratory and wild populations. We used the thoracic fluid of moths from Finland and Georgia as they showed significantly different chemical compositions, and both populations could be successfully reared in the laboratory. The chemical defence fluid from Georgian wood tiger moths reared in the laboratory provoked longer hesitation times compared to the control (coef = -1.91, se = 0.56, z= -3.38, *p*=0.00071; Figure 5A), whereas that of both Finnish laboratory-raised (coef = -0.599, se = 0.46, z= -1.30, p=0.19) and wild moths (coef = -0.50, se = 0.43, z= -1.16, *p*=0.25), and wild Georgian moths (coef = -0.64, se = 0.42, z= -1.51, *p*=0.13) did not differ significantly from the control. When we analyse the amount of oats eaten per unit of time (minutes) which can be used as a proxy for distastefulness, both Finnish (coef = -3.99, se = 0.96, z= -4.17, *p*= 2.98e-05; Figure 5B) and Georgians (coef = -4.10, se = 1.26, z= -3.2, *p*= 0.00119) moth laboratory and Finnish (coef = -2,5, se =0.90, z= -2.78, *p*= 0.00551) and Georgians (coef = -2.15, se = 0.88, z= - 2.43, *p*= 0.01487) wild populations differed from the control (see Figure 5B). We found no significant difference in the beak wiping behaviour between birds exposed to oats soaked in either fluid from laboratory and wild Finnish and Georgian wood tiger moths, and those exposed to water-soaked oats (*p*>0.05, see Figure S4, Table S13, supplementary material). Birds drank more water after tasting the thoracic fluid from laboratory (coef = -3.40, se = 0.94, z= -3.63, *p*= 0.000281) and wild (coef = -3.99, se = 0.94, z= -4.24, *p*= 2.21e-05) Finnish wood tiger moths and wild Georgian wood tiger moths (coef = -2.22, se = 0.86, z= -2.53, *p*= 0.011420, Figure 6).

**Fig. 5.**
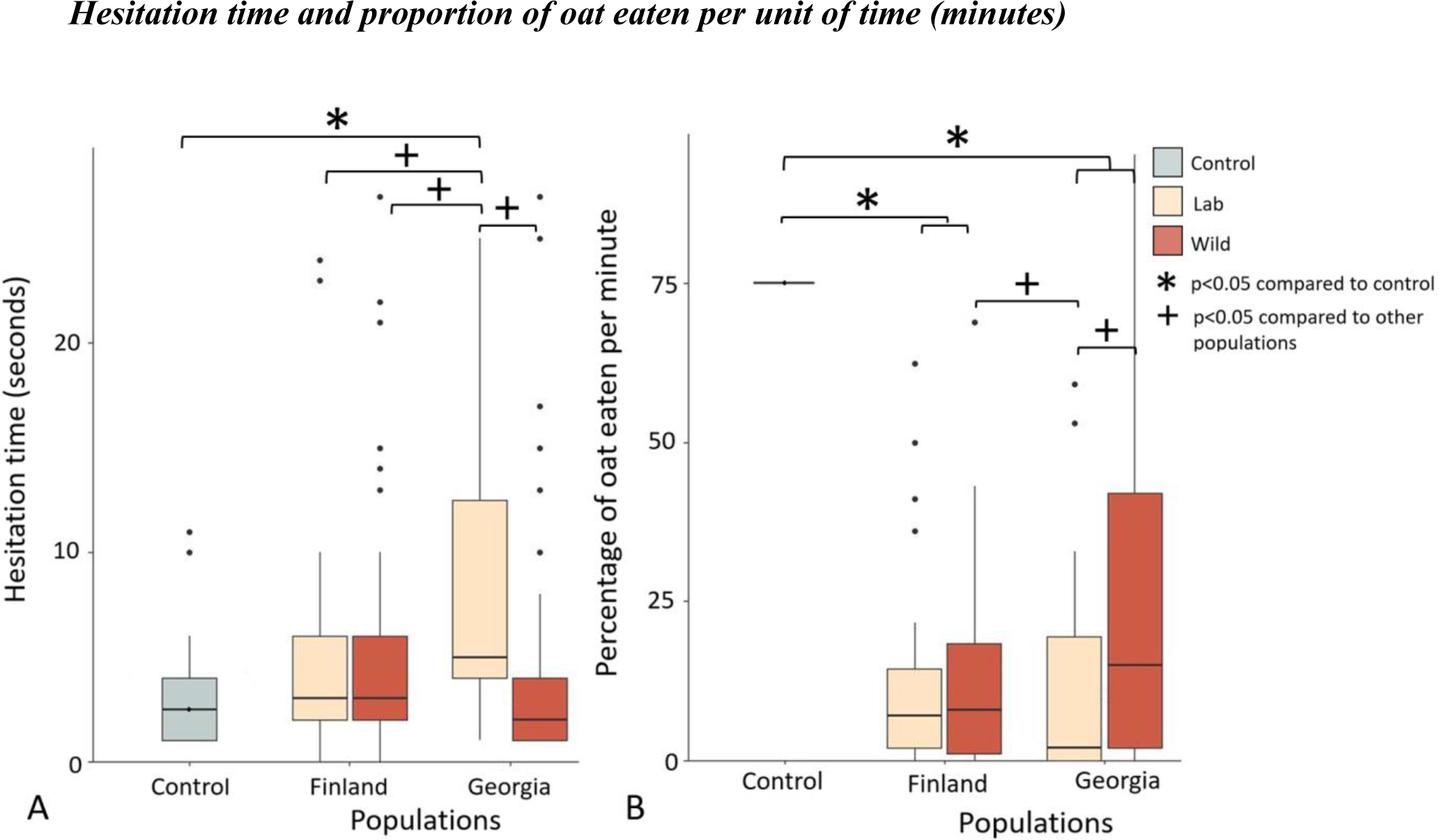
A) Predators hesitate longer when exposed to fluids of Georgia moths raised in laboratory conditions compared to water. Boxes show the median and the 25th and 75th percentiles of data distribution. Vertical lines show the data range. B) Proportion of oat eaten (per unit of time, minutes) in moths’ fluid eaten per minute is lower when predators are exposed to fluids of moths raised in a laboratory and wild condition from both populations compared to water and to the chemical defences of Georgians laboratory moths. Boxes show the median and the 25th and 75th percentiles of data distribution. Vertical lines show the data range.

**Fig. 6.**
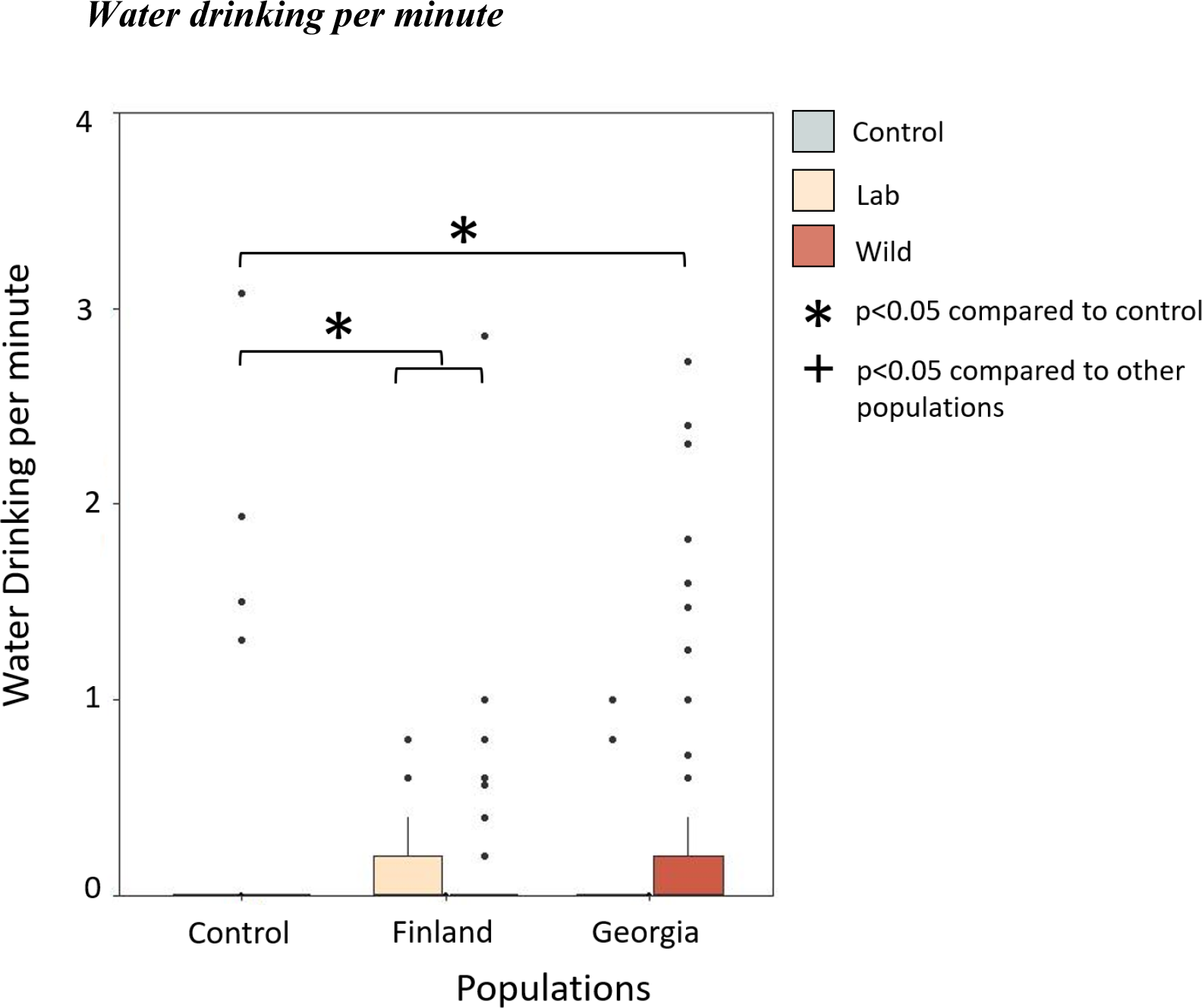
Water drinking rate (per unit of time, minutes) increased when predators are exposed to fluids of moths raised in a laboratory condition from Finnish population and Georgian wild compared to water and to the chemical defences of Georgian moths from laboratory. Water drinking rate not differed between the two populations and wild and laboratory-raised moths. Boxes show the median and the 25th and 75th percentiles of data distribution. Vertical lines show the data range.

The thoracic defence from laboratory-raised Georgian wood tiger moths elicited longer hesitation times in the predators’ response than the defence fluid from Georgian wild moths, and both Finnish laboratory and wild moths (*p*<0.05, see Table S15; supplementary material). The percentage of oat eaten per unit of time (minute), differs between Georgian laboratory moths and Finnish and Georgian wild moths (*p*<0.05, see Table S16; supplementary material). The water drinking behaviour does not differ between Finnish and Georgian population (*p*>0.05, see Table S17; supplementary material).

## 4. Discussion

Insects present an extreme diversification in their antipredator defence strategies. To understand how these defences evolve, and why they vary, it is necessary not only to study the chemical composition of the defences but also to test them on relevant wild predators. In individuals of species with a wide geographic distribution, one may expect a variation in their defences according to characteristics of the environment and the local predation pressure. Predator community composition is likely to cause variation in the defences (Trussell et al., 2000) and diet may indeed play a role, as the diversification in host plants in different regions, for example, can lead to speciation (Nosil et al., 2009). However, the distinctive diversification in the chemical defences in individuals of the same population may not always be explained by the variation in habitat or other external factors: in *Heliconius erato* there is an important genetic component to take into account in the study of intraspecific variation in chemical defences (Mattila et al., 2021).

When looking at the chemical variation across different wood tiger moth populations, we found that the thoracic fluid from laboratory-raised moths had higher amounts of SBMP and IBMP compared to wild-caught moths. The only exception is the Scottish population, where wild individuals had higher levels of SBMP, but not IBMP, compared to individuals raised in the laboratory. Previous studies have shown that *Arctia plantaginis* experienced a different predator community in Scotland, which presented higher predation pressure than that of the other European and Caucasian populations (Nokelainen et al 2014; Rönkä et al 2020). In addition, it has been shown that wild Scottish populations had a higher likelihood of being attacked, as the environment where these moths lived is more open and visible (Nokelainen et al., 2014). This could explain why the Scottish population had higher amounts of SBMP in their defensive fluids and it is in agreement with our findings in this study that birds eat a smaller proportion of oats soaked in high concentrations of pure pyrazine SBMP. The lower variation in IBMP abundance in the wild moths compared to the laboratory sounds counterintuitive at the first glance but since we found that IBMP plays a minor role in chemical defence of moths, the variation seen in the field samples may just reflect the relaxed selection on IBMP.

Notably, the Scottish population was kept only for a few generations in the laboratory, but the Estonian and Finnish populations were established much earlier, albeit with yearly gene flow from the wild in the form of caught individuals. While this might suggest that time in the laboratory may also lead to increased IBMP, it is notable that the Georgian population was also only recently established during the time of sampling and did show a clear difference in IBMP production between the laboratory and the wild. This supports the idea that the greater amounts of pyrazine in the thoracic fluid of *Arctia plantaginis* raised in the laboratory may be due to the constant amount of resources and nutrients that are available in the early life of the individuals, particularly given that interruptions of food availability have been shown to impact their chemical defence (Burdfield-Steel et al. 2019). There are possible trade-offs between the production of pyrazines and other traits that are favoured in the wild moths. Indeed, producing chemical defences *de novo* is costly for *A. plantaginis* (Burdfield-Steel et al. 2018), and deploying those defences is costly too: release of the thoracic fluid decreases the reproductive success in yellow male wood tiger moths (Nokelainen et al., 2012).

Our analysis of wild blue tit responses to pure pyrazines suggests that SBMP alone was a more effective defence than IBMP: birds ate a smaller proportion of oats soaked with SBMP and there was a trend for birds to hesitate longer to approach SBMP oats compared to the control. In contrast, IBMP was a weak defence on its own, although there was a trend for IBMP to cause birds to drink more water, which suggests that birds may find IBMP more aversive after tasting it. Despite having no effect on bird hesitation to approach, the 50/50 blend of IBMP+SBMP influenced the greatest number of bird behaviours: reducing the proportion eaten, increasing drinking behaviour, and there was a trend to increase beak wipes. Rather than the combination having an intermediate effect between that of the two pure pyrazines, as we would expect if the effects of the two were purely additive, this suggests the combination of the two instead has a non-additive, synergistic, effect. The efficacy of this combination during the subjugation stage of attack could explain why moths use IBMP in combination with SBMP even when IBMP alone is mostly ineffective. Similarly, a recent study by Yan et al. (2021) found that sub-threshold pyrazines, which are not detected at the given concentration on their own, can nonetheless contribute synergistically to the organoleptic properties of a chemical mixture as suggested by Maga et al. (1973). Interestingly, Yan et al (2021) also found that sub-threshold pyrazines reduced the odour thresholds of supra-threshold pyrazines, which could explain why the combination of SBMP+IBMP did not affect bird hesitation to approach the defensive odour. Altogether these results suggest that the aversion of a chemical mixture is not the same as the sum of its parts, and chemical defences should therefore be presented in natural combinations to account for potential synergistic and antagonistic relationships that influence the sensory responses of predators.

While SBMP reduced the proportion of oats eaten at the three highest concentrations, surprisingly SBMP+IBMP significantly reduced the proportion eaten at both the lowest and the highest concentrations (0.05 ng/µl, 1.0 ng/µl). Only the 0.5 ng/µl concentration of SBMP+IBMP increased drinking compared to the control and there was a trend for the lowest concentration of IBMP alone (0.05 ng/µl) to increase drinking compared to the control. These results suggest that SBMP is more effective at higher concentrations (by reducing the proportion eaten and increasing hesitation time) and that IBMP is more effective in combination with SBMP and at the lower concentrations (by reducing the proportion eaten and increasing drinking behaviour). Wood tiger moths produce between 0.5 and 2µl of fluid, so the average concentration of the fluids is in the lower range of the concentrations tested (based on the abundances shown in Figure 1). Overall, we found an unexpected relationship with pyrazine concentration, where more is not always better – especially for IBMP. This finding is in line with research on pyrazines in food science, where concentration has been found to change the quality rather than just the intensity of sensory perception. For example, Evers et al. (1972) described 5,7-dihydrothieno (3,4,6) – pyrazine as resembling roasted nuts, baked goods, or fresh milk, depending on the concentration and evaluation medium (Maga et al. 1973). This means that aversion towards a chemical mixture cannot always be extrapolated from the concentration of its contents, and predator responses to defence fluids at natural concentrations should also be measured.

Following the results of the chemical analysis and the wild predator response to pure synthetic pyrazines, we tested the avian predators’ response to the thoracic fluid of Finnish and Georgian *A*.*plantaginis*. The percentage of oats eaten per unit of time was lower for oats soaked in the thoracic fluids of moths from the laboratory. The chemical defences from moths raised in the laboratory from both Finland and Georgia present a more effective defence against the wild blue tit predators by reducing the percentage of oat eaten per minute. This is in line with chemical analyses, which found higher amounts of pyrazines, SBMP and IBMP, in thoracic fluids of moths raised in the laboratory. Variation in the efficacy of chemical defences from individuals of the same population (but raised in different environments, e.g: in wild conditions and laboratory conditions) may be due to food deprivation or competition for the resources (Speed et al., 2012) in wild moths during the early life stages. It has been previously shown (Burdfield-Steel et al., 2019) that resource limitation in early life indeed impacts the efficacy of the wood tiger moth’s chemical defences in terms of bird hesitation time, which was lower when the birds experience the defences of moths raised with fewer nutrients (Burdfield-Steel et al., 2019). Predators, especially birds, can detect the smell of pyrazine from a distance (Guildford et al., 1987), which plays a role in the anti-predator defences of aposematic prey (associated with the warning signal). It has been shown that SBMP (Rojas et al. 2017) and IBMP (Burdfield-Steel et al. 2018) present a strong smell that deters bird attacks. This study found that hesitation time was longer for Georgian moths raised in the laboratory. This may be due to the fact that the Georgian laboratory-raised population was additionally fed with *Plantago sp*. and *Rumex sp* due to the high mortality on *Taraxacum sp* alone (which may in itself suggest different nutritional requirements in this population). Moreover, it is possible that Georgian moths may invest more in the deterrent olfactory cue when they are raised with a constant amount of resources (aka in the laboratory) and on a diet from which they cannot sufficiently sequester defensive toxins such as pyrrolizidine alkaloids (PAs). Predators can indeed use more than one cue to assess the toxicity of prey, so multiple defensive compounds can be used as a multimodal signal (Marples et al., 1994; Rojas et al. 2019). A recent study found that wood tiger moth PAs can also provoke disgust reactions in wild birds (Winters et al., 2021). The presence of PAs alone did not deter the predators, but the combination of both pyrazines and PAs confers better defences to the moths (Winters et al., 2021). Because the laboratory-raised moths in the current study did not sequester PAs from their diet, it is possible that they invested more in the production of pyrazines. Smell is one of the first cues that birds perceive when approaching prey, so this may also explain why laboratory moths seem to allocate more resources to the production of pyrazine.

Finally, it should be noted that wood tiger moths are capital breeders, meaning that adults do not eat, and all resources must be acquired at the larval stage. For that reason, it is unknown how effectively moths can recover their chemical defences after they have released them. Moths can certainly produce defensive fluids multiple times over their lifespan, but the amount of pyrazine may decrease with each release. Evidence from enclosure experiments with wild caught birds and live moths suggests that moths can survive an initial bird attack if rejected up to 30% of the time (Rönka et al. unpublished; Winters et al., 2021). Thus, if wild-caught moths have previously been attacked and released their defensive fluids, this may contribute to the lower level of defence seen. While we cannot rule this out, if prior attacks were indeed driving the pattern of reduced defence in the wild populations, we would expect this to be most noticeable in the Scottish population - where prior studies suggest bird predation is highest and much reduced in the Estonian population where attack rates are low (Rönka et al. 2020). However, this is not the pattern we see (Figure 3), as the Scottish population is in fact the only population not to show this difference between wild and laboratory-reared moths.

Overall, our results suggest that measuring the absolute amounts of chemical defences does not give the full picture of their efficacy: it is also necessary to test chemical defences on relevant predators. The early environment drives variation in methoxypyrazine chemical defences, even though the defences are produced *de novo*. In addition, chemical variation appears to correlate with previously measured predation pressure, suggesting that natural selection may also drive investment in chemical defences in this species. Clearly, the study of chemical defences may be complicated by non-additive interactions between the chemical components of the defence, and caution must be used when extrapolating from chemical measurements to predator responses.

## References

Alonso-Mejia, A. & Brower, L. (1994). From model to mimic: age-dependent unpalatability in monarch butterflies. Cellular and Molecular Life Sciences 50, 176–181.

Arenas, M. L., Walter, D., & Stevens, M. (2015). Signal honesty and predation risk among a closely related group of aposematic species. Scientific Reports, 5(1), 1–12. https://doi.org/10.1038/srep11021

Bartlett, M. S. (1937). Properties of sufficiency and statistical tests. Proceedings of the Royal Society of London. Series A-Mathematical and Physical Sciences, 160(901), 268–282. https://doi.org/10.1098/rspa.1937.0109

Bates, D., Kliegl, R., Vasishth, S., & Baayen, H. (2015). Parsimonious mixed models. arXiv preprint arXiv:1506.04967.

Bezzerides, A. L., McGraw, K. J., Parker, R. S., & Husseini, J. (2007). Elytra color as a signal of chemical defense in the Asian ladybird beetle Harmonia axyridis. Behavioral Ecology and Sociobiology, 61(9), 1401–1408. http://dx.doi.org/10.1007/s00265-007-0371-9

Böttinger, L. C., Hüftlein, F., & Stökl, J. (2021). Mate attraction, chemical defense, and competition avoidance in the parasitoid wasp Leptopilina pacifica. Chemoecology, 31(2), 101–114. https://doi.org/10.1007/s00049-020-00331-3

Bowers, M. D. (1992). The evolution of unpalatability and the cost of chemical defense in insects. In Insect Chemical Ecology: An Evolutionary Approach (eds B. D. Roitberg and M. B. Isman), pp. 216–244. Chapman & Hall, London.

Briolat, E. S., Burdfield-Steel, E. R., Paul, S. C., Rönkä, K. H., Seymoure, B. M., Stankowich, T., & Stuckert, A. M. (2019). Diversity in warning coloration: selective paradox or the norm?. Biological Reviews, 94(2), 388–414. https://doi.org/10.1111/brv.12460

Brower, L., van Brower, J. & Corvino, J. (1967). Plant poisons in a terrestrial food chain. Proceedings of the National Academy of Sciences of the United States of America 57, 893–898. doi:10.1073/pnas.57.4.893

Burdfield-Steel, E., Brain, M., Rojas, B., & Mappes, J. (2019). The price of safety: food deprivation in early life influences the efficacy of chemical defence in an aposematic moth. Oikos, 128(2), 245–253. https://doi.org/10.1111/oik.05420

Burdfield-Steel, E., Pakkanen, H., Rojas, B., Galarza, J. A., & Mappes, J. (2018). De novo synthesis of chemical defenses in an aposematic moth. Journal of Insect Science, 18(2), 28. https://doi.org/10.1093/jisesa/iey020

Cai, L., J. A. Koziel, and M. E. O’Neal. 2007. Determination of characteristic odorants from Harmonia axyridis beetles using in vivo solid-phase microextraction and multidimensional gas chromatography-mass spectrometry-olfactometry. J. Chromatogr. A. 1147: 66–78. doi: 10.1016/j.chroma.2007.02.044.

Chinery, M. 1993. Collins guide to insects of Britain and Western Europe. 3rd ed. Harper-Collins. London.

Cummings, M. E., & Crothers, L. R. (2013). Interacting selection diversifies warning signals in a polytypic frog: an examination with the strawberry poison frog. Evolutionary Ecology, 27(4), 693–710. https://doi.org/10.1007/s10682-013-9648-9

Darst, C. R., & Cummings, M. E. (2006). Predator learning favours mimicry of a less-toxic model in poison frogs. Nature, 440(7081), 208–211. doi: 10.1038/nature04297

Darst, C. R., Cummings, M. E., & Cannatella, D. C. (2006). A mechanism for diversity in warning signals: conspicuousness versus toxicity in poison frogs. Proceedings of the National Academy of Sciences, 103(15), 5852–5857. https://doi.org/10.1073/pnas.0600625103

Eisner, T., Eisner, M. & Siegler, M. (2005). Secret Weapons: Defenses of Insects, Spiders, Scorpions, and Other Many-Legged Creatures. Belknap Press, Harvard. https://doi.org/10.1007/s10841-006-9007-z

Endler, J. A., & Mappes, J. (2004). Predator mixes and the conspicuousness of aposematic signals. The American Naturalist, 163(4), 532–547. doi: 10.1086/382662

Evers, W. J., Wilson, R. A., Theimer, E. T., & Katz, I. (1972). U.S. Patent No. 3, 647,792. Washington, DC: U.S. Patent and Trademark Office.

Guilford, T. I. M., Nicol, C., Rothschild, M., & Moore, B. P. (1987). The biological roles of pyrazines: evidence for a warning odour function. Biological Journal of the Linnean Society, 31(2), 113–128. https://doi.org/10.1111/j.1095-8312.1987.tb01984.x

Ham, A. D., Ihalainen, E., Lindström, L., & Mappes, J. (2006). Does colour matter? The importance of colour in avoidance learning, memorability and generalisation. Behavioral Ecology and Sociobiology, 60(4), 482–491. https://doi.org/10.1007/s00265-006-0190-4

Hegna, R. H., Galarza, J. A., & Mappes, J. (2015). Global phylogeography and geographical variation in warning coloration of the wood tiger moth (Parasemia plantaginis). Journal of Biogeography, 42(8), 1469–1481. https://doi.org/10.1111/jbi.12513

Hudson, C. M., Brown, G. P., Blennerhassett, R. A., & Shine, R. (2021). Variation in size and shape of toxin glands among cane toads from native-range and invasive populations. Scientific Reports, 11(1), 1–11. https://doi.org/10.1038/s41598-020-80191-7

Joron, M., & Iwasa, Y. (2005). The evolution of a Müllerian mimic in a spatially distributed community. Journal of Theoretical Biology, 237(1), 87–103. doi: 10.1016/j.jtbi.2005.04.005

Lawrence J. P., Rojas, B., Fouquet A., Mappes, J., Blanchette A., Saporito R., Bosque R. J., Courtois E., Noonan B. P. (2019) Weak warning signals can persist in the absence of gene flow. PNAS 116:19037–19045. DOI:10.1073/pnas.1901872116.

Leraut, P. (2006). Moths of Europe, Vol. 1: Saturnids, Lasiocampids, Hawkmoths, Tiger Moths… Moths of Europe, Vol. 1: Saturnids, Lasiocampids, Hawkmoths, Tiger Moths…

Lindstedt, C., Talsma, J. H. R., Ihalainen, E., Lindström, L., & Mappes, J. (2010). Diet quality affects warning coloration indirectly: excretion costs in a generalist herbivore. Evolution: International Journal of Organic Evolution, 64(1), 68–78. https://doi.org/10.1111/j.1558-5646.2009.00796.x

Lindstedt, C., Eager, H., Ihalainen, E., Kahilainen, A., Stevens, M., Mappes, J., Direction and strength of selection by predators for the color of the aposematic wood tiger moth, Behavioral Ecology (2011), Pages 580–587, https://doi.org/10.1093/beheco/arr017

Maan, M. E., & Cummings, M. E. (2012). Poison frog colors are honest signals of toxicity, particularly for bird predators. The American Naturalist, 179(1), E1–E14. doi: 10.1086/663197

Maga, J. A., Sizer, C. E., & Myhre, D. V. (1973). Pyrazines in foods. Critical Reviews in Food Science & Nutrition, 4(1), 39–115. https://doi.org/10.1080/10408397309527153

Mallet, J., & Singer, M. C. (1987). Individual selection, kin selection, and the shifting balance in the evolution of warning colours: the evidence from butterflies. Biological Journal of the Linnean Society, 32(4), 337–350. https://doi.org/10.1111/j.1095-8312.1987.tb00435.x

Marples, N. M., van Veelen, W., & Brakefield, P. M. (1994). The relative importance of colour, taste and smell in the protection of an aposematic insect Coccinella septempunctata. Animal Behaviour, 48(4), 967–974. https://doi.org/10.1006/anbe.1994.1322

Mattila, A. L., Jiggins, C. D., Opedal, Ø. H., Montejo-Kovacevich, G., McMillan, W. O., Bacquet, C., & Saastamoinen, M. (2021). Evolutionary and ecological processes influencing chemical defense variation in an aposematic and mimetic Heliconius butterfly. PeerJ, 9, e11523. doi:10.7717/peerj.11523

Moore, B. P., & Brown, W. V. (1981). Identification of warning odour components, bitter principles and antifeedants in an aposematic beetle: Metriorrhynchus rhipidius (Coleoptera: Lycidae). Insect Biochemistry, 11(5), 493–499. https://doi.org/10.1016/0020-1790(81)90016-0

Nokelainen, O., Hegna, R. H., Reudler, J. H., Lindstedt, C., & Mappes, J. (2012). Trade-off between warning signal efficacy and mating success in the wood tiger moth. Proceedings of the Royal Society B: Biological Sciences, 279(1727), 257–265. https://doi.org/10.1098/rspb.2011.0880

Nokelainen, O., Lindstedt, C., & Mappes, J. (2013). Environment-mediated morph-linked immune and life-history responses in the aposematic wood tiger moth. Journal of Animal Ecology, 82(3), 653–662. https://doi.org/10.1111/1365-2656.12037

Nokelainen, O., Valkonen, J., Lindstedt, C., & Mappes, J. (2014). Changes in predator community structure shifts the efficacy of two warning signals in Arctiid moths. Journal of Animal Ecology, 598–605. https://doi.org/10.1111/1365-2656.12169

Nosil, P., Harmon, L. J., & Seehausen, O. (2009). Ecological explanations for (incomplete) speciation. Trends in ecology & evolution, 24(3), 145–156. doi: 10.1016/j.tree.2008.10.011

Ojala, K., Julkunen-Tiitto, R., Lindström, L., & Mappes, J. (2005). Diet affects the immune defence and life-history traits of an Arctiid moth Parasemia plantaginis. Evolutionary Ecology Research, 7(8), 1153–1170.

Pfeiffer, L., Ruther, J., Hofferberth, J., & Stökl, J. (2018). Interference of chemical defence and sexual communication can shape the evolution of chemical signals. Scientific Reports, 8(1), 1–10. https://doi.org/10.1038/s41598-017-18376-w

Powell, J. A., & Opler, P. A. (2009). Moths of western north america. In Moths of Western North America. University of California Press.

Rojas, B., E. Burdfield-Steel, H. Pakkanen, K. Suisto, M. Maczka, S. Schulz, and J. Mappes. (2017). How to fight multiple enemies: target-specific chemical defences in an aposematic moth. Proc. R. Soc. B 284: 20171424. https://doi.org/10.1098/rspb.2017.1424

Rojas B., Mappes J., Burdfield-Steel E. (2019). “Multiple modalities in insect warning displays have additive effects against wild avian predators.” Behavioral Ecology and Sociobiology 73(3): 37. https://doi.org/10.1007/s00265-019-2643-6

Rojas B., Valkonen J.K., Nokelainen O. (2015) Aposematism. Current Biology 25: R350–R351 https://doi.org/10.1016/j.cub.2015.02.015

Rönkä, K., J. Mappes, L. Kaila, and N. Wahlberg. (2016). Putting Parasemia in its phylogenetic place: a molecular analysis of the subtribe Arctiina (Lepidoptera). Syst. Entomol. 41: 844–853. https://doi.org/10.1111/syen.12194

Rönkä, K., De Pasqual, C., Mappes, J., Gordon, S., & Rojas, B. (2018). Colour alone matters: no predator generalization among morphs of an aposematic moth. Animal Behaviour, 135, 153–163. DOI: 10.1016/j.anbehav.2017.11.015

Rönkä, K., Valkonen, J. K., Nokelainen, O., Rojas, B., Gordon, S., Burdfield-Steel, E., & Mappes, J. (2020). Geographic mosaic of selection by avian predators on hindwing warning colour in a polymorphic aposematic moth. Ecology Letters, 23(11), 1654–1663.

Rothschild, M., & Moore, B. (1987). Pyrazines as alerting signals in toxic plants and insects. Insects-Plants. Dordrecht, Holland, 97–107. https://doi.org/10.1111/ele.13597

Rowe, C., & Guilford, T. (1996). Hidden colour aversions in domestic chicks triggered by pyrazine odours of insect warning displays. Nature, 383(6600), 520–522.

Ruxton, G., Sherratt, T. & Speed, M. (2004). Avoiding Attack: The Evolutionary Ecology of Crypsis, Warning Signals, and Mimicry. Oxford University Press, Oxford, UK. DOI:10.1093/acprof:oso/9780198528609.001.0001

Ruxton, Graeme D., William L. Allen, Thomas N. Sherratt, and Michael P. Speed. Avoiding Attack: The Evolutionary Ecology of Crypsis, Aposematism, and Mimicry. 2nd ed. Oxford: Oxford University Press, 2018. Oxford Scholarship Online, 2018. doi:10.1093/oso/9780199688678.001.0001.

Sculfort, O., de Castro, E. C., Kozak, K. M., Bak, S., Elias, M., Nay, B., & Llaurens, V. (2020). Variation of chemical compounds in wild Heliconiini reveals ecological factors involved in the evolution of chemical defenses in mimetic butterflies. Ecology and Evolution, 10(5), 2677–2694. doi:10.1002/ece3.6044

Sherratt, T. N., Speed, M. P., & Ruxton, G. D. (2004). Natural selection on unpalatable species imposed by state-dependent foraging behaviour. Journal of Theoretical Biology, 228(2), 217–226. doi: 10.1016/j.jtbi.2003.12.009

Sherratt, T. N. (2006). Spatial mosaic formation through frequency-dependent selection in Müllerian mimicry complexes. Journal of theoretical biology, 240(2), 165–174. doi: 10.1016/j.jtbi.2005.09.017

Sherratt, T. N. (2008). The evolution of Müllerian mimicry. Naturwissenschaften, 95(8), 681–695. doi: 10.1007/s00114-008-0403-y

Speed M. P., Ruxton G. D., Mappes J. (2012) Why are defensive toxins so variable? An evolutionary perspective. Biological Reviews. Volume 87, Issue 4. https://doi.org/10.1111/j.1469-185X.2012.00228.x

Therneau, T. 2020. Coxme: Mixed Effects Cox Models. http://CRAN.R-project.org/package=coxme.

Trussell, G. C., & Smith, L. D. (2000). Induced defenses in response to an invading crab predator: an explanation of historical and geographic phenotypic change. Proceedings of the National Academy of Sciences, 97(5), 2123–2127. https://doi.org/10.1073/pnas.040423397

Watson, A., & Goodger, D. T. (1986). Catalogue of the Neotropical tiger-moths. British Museum (Natural History).

Weldon, P. J. (2017). Poison frogs, defensive alkaloids, and sleepless mice: critique of a toxicity bioassay. Chemoecology, 27(4), 123–126. https://doi.org/10.1007/s00049-017-0238-0

Winters, A. E., White, A. M., Cheney, K. L., & Garson, M. J. (2019). Geographic variation in diterpene-based secondary metabolites and level of defence in an aposematic nudibranch, Goniobranchus splendidus. Journal of Molluscan Studies, 85(1), 133–142. https://doi.org/10.1111/1365-2656.13643

Winters, A. E., Lommi, J., Kirvesoja, J., Nokelainen, O., & Mappes, J. (2021). Multimodal aposematic defenses through the predation sequence. Frontiers in Ecology and Evolution, 512. https://doi.org/10.3389/fevo.2021.657740

Yan, Y., Chen, S., Nie, Y., & Xu, Y. (2021). Quantitative Analysis of Pyrazines and Their Perceptual Interactions in Soy Sauce Aroma Type Baijiu. Foods, 10(2), 441. https://doi.org/10.3390/foods10020441

Yen, E. C., McCarthy, S. A., Galarza, J. A., Generalovic, T. N., Pelan, S., Nguyen, P., … & Jiggins, C. D. (2020). A haplotype-resolved, de novo genome assembly for the wood tiger moth (Arctia plantaginis) through trio binning. GigaScience, 9(8), giaa088. https://doi.org/10.1093/gigascience/giaa088

