## Supplementary material for "Not just the sum of its parts: geographic variation and non-additive effects of pyrazines in the chemical defence of an aposematic moth"

**Title:** Geographic variation in wood tiger moth (*Arctia plantaginis*) chemical defence has non-additive effects against birds

Cristina Ottocento1,4, Anne E. Winters1,2, Bibiana Rojas1,3, Johanna Mappes1,4, and Emily Burdfield-Steel1,5

1University of Jyväskylä, Department of Biology and Environmental Science, P.O. Box 35, 40014, Jyväskylä, Finland

2University of Exeter, College Life and Environmental Sciences, Penryn Campus, Penryn, TR10 9FE, United Kingdom

3 Department of Interdisciplinary Life Sciences, Konrad Lorenz Institute of Ethology, University of Veterinary Medicine Vienna, Savoyenstraße 1, 1160, Vienna, Austria

4University of Helsinki, Organismal and Evolutionary Biology Research Programme, Faculty of Biological and Environmental Sciences, P.O. Box 65, Viikki Biocenter 3, Finland

5University of Amsterdam, Institute for Biodiversity and Ecosystem Dynamics, Science Park 904, 1098 XH Amsterdam, The Netherlands

**Contents:	Pages**

**Differences in pyrazine levels across populations**	2-3

Table S1, table S2, table S3, table S4, table S5, table S6

**Bird response to pure pyrazine treatments**	4-9

Figure S1, table S7, table S8, figure S2,

Table S9, figure S3, table S10

**Bird response to moths’ defensive fluid**	9-14

Table S11, table S12, figure S4, table S13,

Table S14, table S15, table S16, table S17

**Differences in pyrazine levels across populations**

**TableS1**. The amount of SBMP in laboratory and wild populations of wood tiger moths was analysed using linear mixed-effects model with a normal distribution. In the model, SBMP variation was set as dependent variable and the different populations, the origin and the interactions between populations and origin as explanatory variable. Each model also included year as a random factor.

|  | Chisq | Df | Pr(>Chisq) |
| --- | --- | --- | --- |
| population | 5.4246 | 3 | 0.14322 |
| origin (laboratory/wild) | 0.1102 | 1 | 0.73991 |
| population:origin | 10.6759 | 3 | 0.01361 * |

**TableS2.** The amount of IBMP in laboratory and wild populations of wood tiger moths was analysed using linear mixed-effects model with a normal distribution. In the model, IBMP variation was set as dependent variable and the different populations, the origin and the interactions between populations and origin as explanatory variable. Each model also included year as a random factor.

|  | Chisq | Df | Pr(>Chisq) |
| --- | --- | --- | --- |
| population | 27.2475 | 3 | 5.224e-06 *** |
| origin (laboratory/wild) | 11.0824 | 1 | 0.0008715 *** |
| population:origin | 5.7327 | 3 | 0.1253663 |

**TableS3.** Variance in SBMP and IBMP in wild and laboratory wood tiger moth populations

|  | variance WILD | variance LAB |
| --- | --- | --- |
| SBMP | 0.02470923 | 0.02453391 |
| IBMP | 0.008763562 | 0.023683064 |

**TableS4.** Variance in SBMP and IBMP in Estonia, Finland, Georgia and Scotland wood tiger moth populations

|  | variance Estonians | variance Finnish | variance Georgians | variance Scottish |
| --- | --- | --- | --- | --- |
| SBMP | 0.01300987 | 0.03052636 | 0.01509857 | 0.01939160 |
| IBMP | 0.018332771 | 0.017158750 | 0.003576965 | 0.016252634 |

**TableS5.** The ratio of IBMP to SBMP in laboratory and wild populations of wood tiger moths was analysed using linear mixed-effects model with a normal distribution. The ratio was set as dependent variable and the different populations, the origin and the interactions between populations and origin as explanatory variable, year was included as a random factor.

|  | Chisq | Df | Pr(>Chisq) |
| --- | --- | --- | --- |
| population | 18.0482 | 3 | 0.0004299 *** |
| origin (laboratory/wild) | 2.6254 | 1 | 0.1051681 |
| population:origin | 1.9633 | 3 | 0.5800483 |

**TableS6.** The total amount of pyrazine in laboratory and wild populations of wood tiger moths was analysed using linear mixed-effects model with a normal distribution. The total amount of pyrazine was set as dependent variable and the different populations, the origin and the interactions between populations and origin as explanatory variable, year was included as a random factor.

|  | Chisq | Df | Pr(>Chisq) |
| --- | --- | --- | --- |
| population | 12.5799 | 3 | 0.005639 ** |
| origin(laboratory/wild) | 4.7878 | 1 | 0.028663 * |
| population:origin | 10.2273 | 3 | 0.016730 * |

**Bird response to pure pyrazine treatments**

**Hesitation time compared to control (water)**

**FigS1.** Bird hesitation to approach fluid-soaked oats (per second) for each pyrazine type (SBMP = red, IBMP = yellow, SBMP+IBMP = orange) and ng/ul concentration (higher concentrations shown in darkening shades) compared to the water control. Boxes show the median and the 25th and 75th percentiles of data distribution. Vertical lines show the data range.

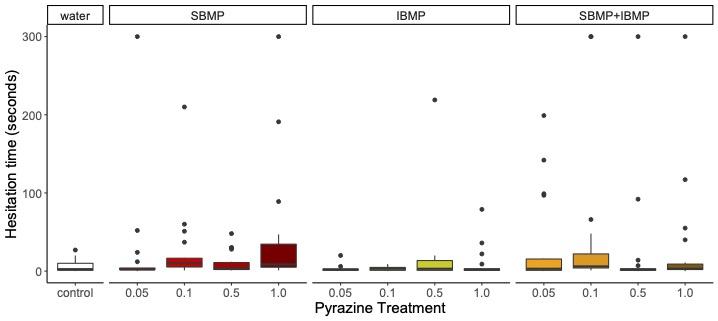

**TableS7.** Hesitation time was analyzed using a cox proportional hazards model. Hesitation time was set as the dependent variable with chemical treatment and trial number as explanatory variables. Trial duration was included as an offset term and bird ID was included as a random factor.

|  | coef | exp(coef) | se(coef) | z | p |
| --- | --- | --- | --- | --- | --- |
| 0.05.SBMP | 0.02110384 | 1.0213281 | 0.69651554 | 0.03 | 0.980 |
| 0.1.SBMP | -1.33234319 | 0.2638583 | 0.68206040 | -1.95 | 0.051 |
| 0.5.SBMP | -0.51523720 | 0.5973589 | 0.68376214 | -0.75 | 0.450 |
| 1.SBMP | -1.33568819 | 0.2629771 | 0.68597538 | -1.95 | 0.052 |
| 0.05.IBMP | 0.57665048 | 1.7800661 | 0.67829238 | 0.85 | 0.400 |
| 0.1.IBMP | 0.24284442 | 1.2748703 | 0.68077454 | 0.36 | 0.720 |
| 0.5.IBMP | -0.32925586 | 0.7194589 | 0.72447363 | -0.45 | 0.650 |
| 1.IBMP | -0.09995016 | 0.9048825 | 0.68332320 | -0.15 | 0.880 |
| 0.05.SBMP+IBMP | -0.44034420 | 0.6438148 | 0.68749887 | -0.64 | 0.520 |
| 0.1.SBMP+IBMP | -1.07497975 | 0.3413047 | 0.69152288 | -1.55 | 0.120 |
| 0.5.SBMP+IBMP | -0.30085112 | 0.7401880 | 0.68383166 | -0.44 | 0.660 |
| 1.SBMP+IBMP | -0.59416081 | 0.5520256 | 0.63982850 | -0.93 | 0.350 |
| trial | -0.21179495 | 0.8091306 | 0.09297766 | -2.28 | 0.023 |

**Percentage of oats eaten per minute compared to control (water)**

**TableS8**. The proportion of oats eaten per minute was analyzed using a Generalized linear mixed model. The proportion of oats eaten was set as the dependent variable with chemical treatment and trial number as explanatory variables. Trial duration was included as an offset term and bird ID was included as a random factor.

|  | Estimate | Std. Error | z-value | Pr(>\|z\|) |
| --- | --- | --- | --- | --- |
| Intercept | 1.919857 | 0.907639 | 2.115 | 0.034411 * |
| 0.05.SBMP | -1.73342 | 1.28409 | -1.35 | 0.177041 |
| 0.1.SBMP | -2.702417 | 1.283778 | -2.105 | 0.035287 * |
| 0.5.SBMP | -4.756727 | 1.308055 | -3.636 | 0.000276 *** |
| 1.SBMP | -3.00573 | 1.283983 | -2.341 | 0.019235 * |
| 0.05.IBMP | -1.531945 | 1.283504 | -1.194 | 0.232648 |
| 0.1.IBMP | -2.105292 | 1.284191 | -1.639 | 0.101132 |
| 0.5.IBMP | -0.948057 | 1.346029 | -0.704 | 0.481223 |
| 1.IBMP | 0.003231 | 1.283642 | 0.003 | 0.997992 |
| 0.05.SBMP+IBMP | -2.568568 | 1.28494 | -1.999 | 0.045611 * |
| 0.1.SBMP+IBMP | -1.926902 | 1.286213 | -1.498 | 0.134102 |
| 0.5.SBMP+IBMP | -1.640855 | 1.28314 | -1.279 | 0.200974 |
| 1.SBMP+IBMP | -2.350563 | 1.200908 | -1.957 | 0.050310 . |
| trial | -0.066418 | 0.011857 | -5.602 | 2.12e-08 *** |

**Beak wipes per minute compared to control (water)**

**FigS2.** Number of beak wipes per minute after eating fluid-soaked oats for each pyrazine type (SBMP = red, IBMP = yellow, SBMP+IBMP = orange) and ng/ul concentration (higher concentrations shown in darkening shades) compared to the water control. Boxes show the median and the 25th and 75th percentiles of data distribution. Vertical lines show the data range.

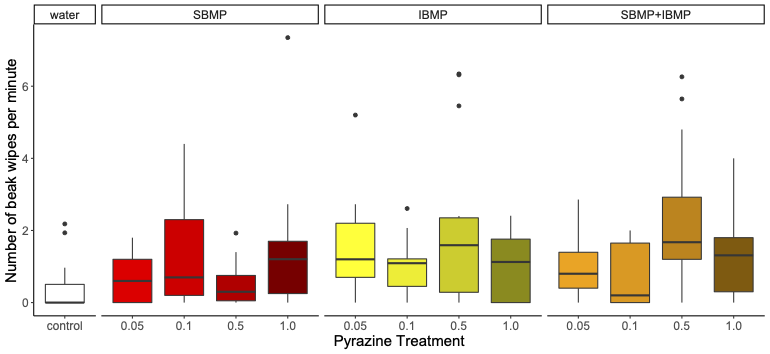

**TableS9.** The number of beak wipes per minute was analyzed using a Generalized linear mixed model. The number of beak wipes was set as the dependent variable with chemical treatment and trial number as explanatory variables. Trial duration was included as an offset term and bird ID was included as a random factor.

|  | Estimate | Std. Error | z-value | Pr(>\|z\|) |
| --- | --- | --- | --- | --- |
| Intercept | -3.86983 | 0.65043 | -5.95 | 2.69E-09 *** |
| 0.05.SBMP | 0.35456 | 0.8576 | 0.413 | 0.6793 |
| 0.1.SBMP | 0.21638 | 0.83181 | 0.26 | 0.7948 |
| 0.5.SBMP | -0.36344 | 0.92809 | -0.392 | 0.6954 |
| 1.SBMP | 0.45822 | 0.82543 | 0.555 | 0.5788 |
| 0.05.IBMP | 1.32915 | 0.83182 | 1.598 | 0.1101 |
| 0.1.IBMP | 0.96899 | 0.82049 | 1.181 | 0.2376 |
| 0.5.IBMP | 1.39332 | 0.85733 | 1.625 | 0.1041 |
| 1.IBMP | 1.08663 | 0.87526 | 1.241 | 0.2144 |
| 0.05.SBMP+IBMP | 0.79357 | 0.83485 | 0.951 | 0.3418 |
| 0.1.SBMP+IBMP | -0.21947 | 0.91336 | -0.24 | 0.8101 |
| 0.5.SBMP+IBMP | 1.57634 | 0.81862 | 1.926 | 0.0542 . |
| 1.SBMP+IBMP | 0.61182 | 0.79468 | 0.77 | 0.4414 |
| trial | -0.01961 | 0.04572 | -0.429 | 0.6681 |

**Water drinks per minute compared to control (water)**

**FigS3.** Number of water drinks per minute after eating fluid-soaked oats for each pyrazine type (SBMP = red, IBMP = yellow, SBMP+IBMP = orange) and ng/ul concentration (higher concentrations shown in darkening shades) compared to the water control. Boxes show the median and the 25th and 75th percentiles of data distribution. Vertical lines show the data range.

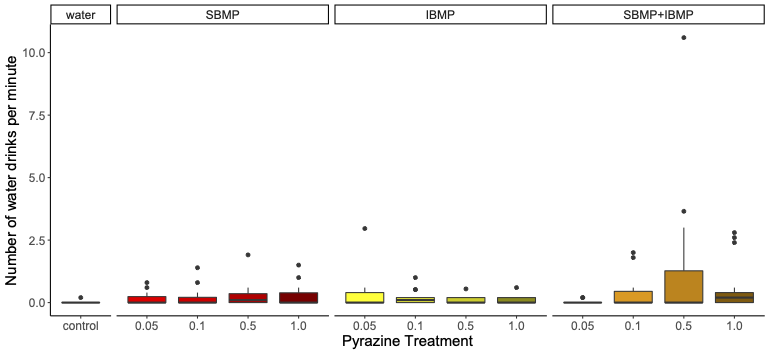

**TableS10.** The number of water drinks per minute was analyzed using a Generalized linear mixed model. The number of water drinks was set as the dependent variable with chemical treatment and trial number as explanatory variables. Trial duration was included as an offset term and bird ID was included as a random factor.

|  | Estimate | Std. Error | z-value | Pr(>\|z\|) |
| --- | --- | --- | --- | --- |
| Intercept | -7.91351 | 1.29083 | -6.131 | 8.76E-10 *** |
| 0.05.SBMP | 1.8867 | 1.44936 | 1.302 | 0.1930 |
| 0.1.SBMP | 1.43819 | 1.41665 | 1.015 | 0.3100 |
| 0.5.SBMP | 2.09028 | 1.47166 | 1.42 | 0.1555 |
| 1.SBMP | 1.79068 | 1.42042 | 1.261 | 0.2074 |
| 0.05.IBMP | 2.45949 | 1.41794 | 1.735 | 0.0828 . |
| 0.1.IBMP | 1.78165 | 1.43079 | 1.245 | 0.2131 |
| 0.5.IBMP | 1.71286 | 1.48636 | 1.152 | 0.2492 |
| 1.IBMP | 1.55133 | 1.51563 | 1.024 | 0.3060 |
| 0.05.SBMP+IBMP | 0.35364 | 1.60208 | 0.221 | 0.8253 |
| 0.1.SBMP+IBMP | 2.34631 | 1.46164 | 1.605 | 0.1084 |
| 0.5.SBMP+IBMP | 3.05731 | 1.41173 | 2.166 | 0.0303 * |
| 1.SBMP+IBMP | 1.97975 | 1.38978 | 1.425 | 0.1543 |
| trial | 0.33872 | 0.07357 | 4.604 | 4.15E-06 *** |

**Bird response to moths’ defensive fluid**

**Table S11.** Hesitation time (in seconds) with control, water. To test the difference in hesitation time in response to thoracic fluids from wild and lab Finnish and Georgian wood tiger moths, we used a cox proportional hazards model using the package coxme. The model included bird ID as a random factor. The population, origin (wild/laboratory) and the interaction between the two were set as fixed factors. Also, the trial with duration as an offset was included as an explanatory variable, while the hesitation time was set as response variables.

|  | coef | exp(coef) | se(coef) | z | p |
| --- | --- | --- | --- | --- | --- |
| Finnish laboratory | -0.59959568 | 0.5490336 | 0.4606419 | -1.30 | 0.19000 |
| Georgian laboratory | -1.90984069 | 0.1481040 | 0.5643646 | -3.38 | 0.00071 |
| Finnish wild | -0.49860078 | 0.6073799 | 0.4313979 | -1.16 | 0.25000 |
| Georgian wild | -0.63693985 | 0.5289085 | 0.4207335 | -1.51 | 0.13000 |
| Trial 3 | -0.01873599 | 0.9814384 | 0.1460923 | -0.13 | 0.90000 |

**TableS12.** Percentage of oat eaten per minute with control (water). The percentage of oats eaten per minute was tested using general linear mixed-effects model with Poisson distribution. The model included bird ID as a random factor. The population, origin (wild/laboratory) and the interaction between the two were set as fixed factors. Also, the trial with duration as an offset was included as an explanatory variable, while the percentage of oat eaten was set as response variables.

|  | Estimate | Std. Error | z-value | Pr(>\|z\|) |
| --- | --- | --- | --- | --- |
| Intercept | 3.96565 | 0.79956 | 4.960 | 7.06e-07 *** |
| Finnish laboratory | -3.99115 | 0.95592 | -4.175 | 2.98e-05 *** |
| Georgian laboratory | -4.10233 | 1.26536 | -3.242 | 0.00119 ** |
| Finnish wild | -2.50389 | 0.90210 | -2.776 | 0.00551 ** |
| Georgian wild | -2.14593 | 0.88113 | -2.435 | 0.01487 * |
| Trial 3 | 0.16099 | 0.01837 | 8.763 | < 2e-16 *** |

**Fig S4.** Beak cleaning (per minute) with control, water. There are no differences in the beak cleaning behaviour when predators are exposed to fluids of moths raised in a laboratory and wild condition from both populations compared to water. Boxes show the median and the 25th and 75th percentiles of data distribution. Vertical lines show the data range

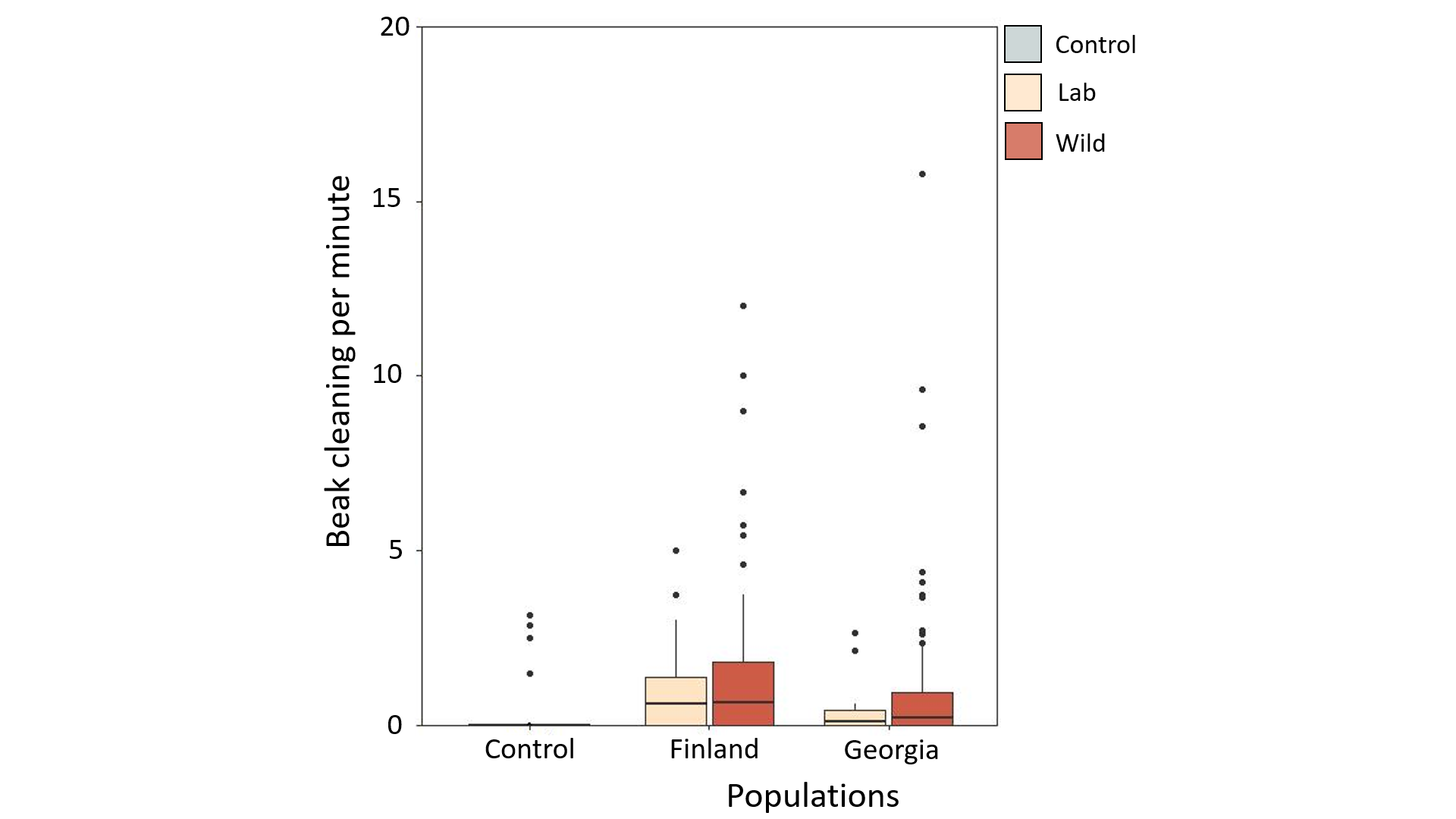

**Table S13.** Beak cleaning per minute with control (water). The beak cleaning was tested using general linear mixed-effects model with Poisson distribution. The model included bird ID as a random factor. The population, origin (wild/laboratory) and the interaction between the two were set as fixed factors. Also, the trial with duration as an offset was included as an explanatory variable, while the beak cleaning per minute was set as response variables.

|  | Estimate | Std. Error | z-value | Pr(>\|z\|) |
| --- | --- | --- | --- | --- |
| Intercept | -2.84303 | 0.79506 | -3.576 | 0.000349 *** |
| Finnish laboratory | -0.64921 | 0.88178 | -0.736 | 0.461579 |
| Georgian laboratory | -0.87944 | 1.07065 | -0.821 | 0.411412 |
| Finnish wild | 0.01733 | 0.85456 | 0.020 | 0.983821 |
| Georgian wild | -0.04596 | 0.84017 | -0.055 | 0.956376 |

**Table S14.** Water drinking per minute with control (water). The water drinking was tested using general linear mixed-effects model with Poisson distribution. The model included bird ID as a random factor. The population, origin (wild/laboratory) and the interaction between the two were set as fixed factors. Also, the trial with duration as an offset was included as an explanatory variable, while the water drinking per minute was set as response variables.

|  | Estimate | Std. Error | z-value | Pr(>\|z\|) |
| --- | --- | --- | --- | --- |
| Intercept | -2.14891 | 0.80594 | -2.666 | 0.007668 ** |
| Finnish laboratory | -3.40204 | 0.93657 | -3.632 | 0.000281 *** |
| Georgian laboratory | -22.72905 | 512.00040 | -0.044 | 0.964591 |
| Finnish wild | -3.98899 | 0.94017 | -4.243 | 2.21e-05 *** |
| Georgian wild | -2.18547 | 0.86397 | -2.530 | 0.011420 * |
| Trial 3 | 0.03443 | 0.20988 | 0.164 | 0.869691 |

**TableS15.** Hesitation time without control. We compared differences between populations using Tukey–Kramer post hoc test for multiple comparisons and excluding the water control group. Statistical significance was set at p < 0.05.

|  | diff | lwr | upr | p adj |
| --- | --- | --- | --- | --- |
| geo.lab-fin.lab | 49.279762 | 4.988939 | 93.570585 | 0.0225133 |
| fin.wild-fin.lab | -13.036415 | -42.623462 | 16.550633 | 0.6645474 |
| geo.wild-fin.lab | 4.291126 | -23.983059 | 32.565310 | 0.9793555 |
| fin.wild-geo.lab | -62.316176 | -104.206101 | -20.426251 | 0.0008883 |
| geo.wild-geo.lab | -44.988636 | -85.961822 | -4.015451 | 0.0250930 |
| geo.wild-fin.wild | 17.327540 | -7.014159 | 41.669239 | 0.2560144 |

**Table S16.** Percentage of oat eaten per minute without control. We compared differences between populations using Tukey–Kramer post hoc test for multiple comparisons and excluding the water control group. Statistical significance was set at p < 0.05.

|  | diff | lwr | upr | p adj |
| --- | --- | --- | --- | --- |
| geo.lab-fin.lab | -15.514881 | -46.530357 | 15.50059 | 0.5668428 |
| fin.wild-fin.lab | 15.738796 | -4.980087 | 36.45768 | 0.2037713 |
| geo.wild-fin.lab | 14.672619 | -5.126906 | 34.47214 | 0.2231397 |
| fin.wild-geo.lab | 31.253676 | 1.919474 | 60.58788 | 0.0317807 |
| geo.wild-geo.lab | 30.187500 | 1.495262 | 58.87974 | 0.0349340 |
| geo.wild-fin.wild | -1.066176 | -18.111906 | 15.97955 | 0.9984832 |

**Table S17.** Water drinking per minute without control. We compared differences between populations using Tukey–Kramer post hoc test for multiple comparisons and excluding the water control group. Statistical significance was set at p < 0.05.

|  | diff | lwr | upr | p adj |
| --- | --- | --- | --- | --- |
| geo.lab-fin.lab | -0.42559524 | -1.5361760 | 0.6849856 | 0.7539066 |
| fin.wild-fin.lab | -0.34103641 | -1.0829239 | 0.4008511 | 0.6335521 |
| geo.wild-fin.lab | 0.13690476 | -0.5720630 | 0.8458725 | 0.9589684 |
| fin.wild-geo.lab | 0.08455882 | -0.9658201 | 1.1349377 | 0.9967872 |
| geo.wild-geo.lab | 0.56250000 | -0.4648919 | 1.5898919 | 0.4896008 |
| geo.wild-fin.wild | 0.47794118 | -0.1324206 | 1.0883029 | 0.1810604 |
